# Bacteriophage Resistance Affects *Flavobacterium columnare* Virulence Partly via Mutations in Genes Related to Gliding Motility and Type IX Secretion System

**DOI:** 10.1101/2020.10.02.323337

**Authors:** Heidi M. T. Kunttu, Anniina Runtuvuori-Salmela, Krister Sundell, Tom Wiklund, Mathias Middelboe, Lotta Landor, Roghaieh Ashrafi, Ville Hoikkala, Lotta-Riina Sundberg

**Author notes:** Address correspondence to Heidi M. T. Kunttu. Lotta Landor, Department of Biological Sciences, University of Bergen, Bergen, Norway.

## Abstract

Increasing problems with antibiotic resistance has directed interest towards phages as tools to treat bacterial infections in the aquaculture industry. However, phage resistance evolves rapidly in bacteria posing a challenge for successful phage therapy. To investigate phage resistance in the fish pathogenic bacterium *Flavobacterium columnare*, two phage-sensitive, virulent wild-type isolates, FCO-F2 and FCO-F9, were exposed to phages and subsequently analyzed for bacterial viability and colony morphology. Twenty-four phage-exposed isolates were further characterized for phage resistance, antibiotic susceptibility, motility, adhesion and biofilm formation on polystyrene surface, protease activity, whole genome sequencing and virulence against rainbow trout fry. Bacterial viability first decreased in the exposure cultures, subsequently increasing after 1-2 days. Simultaneously, the colony morphology of the phage-exposed isolates changed from original rhizoid to rough. The rough isolates arising in phage exposure were phage-resistant with low virulence, whereas rhizoid isolates maintained phage sensitivity, though reduced, and high virulence. Gliding motility and protease activity were also related to the phage sensitivity. Observed genetic mutations in phage-resistant isolates were mostly located in genes coding for type IX secretion system, a component of the flavobacterial gliding motility machinery. However, there were mutational differences between individual isolates, and not all phage-resistant isolates had genetic mutations. This indicates that development of phage resistance in *F. columnare* probably is a multifactorial process including both genetic mutations and changes in gene expression. Phage resistance may not, however, be a challenge for development of phage therapy against *F. columnare* infections, since phage resistance is associated with decrease in bacterial virulence.

**Importance:** Phage resistance of infectious bacteria is a common phenomenon posing challenges for development of phage therapy. Along with growing World population and need for increased food production, constantly intensifying animal farming has to face increasing problems of infectious diseases. Columnaris disease, caused by *F. columnare*, is a worldwide threat for salmonid fry and juvenile farming. Without antibiotic treatments, infections can lead to 100% mortality in a fish stock. Phage therapy of columnaris disease would reduce a development of antibiotic-resistant bacteria and antibiotic loads by the aquaculture industry, but phage-resistant bacterial isolates may become a risk. However, phenotypic and genetic characterization of phage-resistant *F. columnare* isolates in this study revealed that they are less virulent than phage-sensitive isolates and thus not a challenge for phage therapy against columnaris disease. This is a valuable information for the fish farming industry globally when considering phage-based prevention and curing methods for *F. columnare* infections.

## Introduction

Aquaculture has a central role in supporting the increasing demand for high quality protein and healthy food. However, the use of chemotherapy in disease treatment in the industry has led to increased resistance of disease-causing agents to commonly used antibiotics (1, 2). Further, in the face of climate warming, production of protein with smaller carbon footprint is of increasing importance. This has put a pressure on aquaculture industry to increase efficiency in food production, which also means developing more effective ways to fight infectious diseases in intensive farming including reduction the use of antibiotics. Although vaccines against many microbial diseases are in use globally in aquaculture, there are still many diseases with no potent immunization method available (3). This applies especially to infections of fish fry, where efficiency of vaccination is poor due to lack of development of fish secondary immunity at the early life stage.

One of these diseases affecting fry is caused by the fish pathogenic bacterium *Flavobacterium columnare*, the infectious agent of columnaris disease. Columnaris infections cause extensive losses in farmed salmonid fry and juveniles, populations of different catfish species and ayu (*Plecoglossus altivelis*) around the world in water temperatures above 18 °C. The only effective curing method is antibiotic treatment. However, infections often occur repeatedly and may cause up to 100% mortality in rainbow trout fry populations if not treated, thus causing major economic losses to the industry (4, 5). In addition, elevated water temperatures due to warmer summers in the recent years are suggested to enhance virulence development in *F. columnar*e (5). Although antibiotic resistance in this bacterium is not yet as severe problem as in related pathogens, e.g. *Flavobacterium psychrophilum* (6, 7) or *Vibrio* species (8, 9), strains that have acquired resistance towards commonly used antibiotics already exist (10).

Bacteriophages (phages) are viruses that specifically infect their host bacteria, without harming the surrounding microbial community (reviewed in 11). Among the alternatives to traditional antibiotics, phage therapy, i.e. the use of phages against bacterial infections, has demonstrated a strong potential for controlling disease outbreaks in aquaculture (e.g. 12-14). Promising results have been gained also in phage therapy trials of Flavobacterial infections. In a study by Castillo et al. (15), phage treatment reduced the mortality of *F. psychrophilum*-infected Atlantic salmon (*Salmo salar*) by 60 % and rainbow trout (*Oncorhynchus mykiss*) by 67 %. In studies with columnaris infections, mortality of zebra fish (*Danio rerio*) and rainbow trout were reduced by 100 % and nearly 42 %, respectively, in the presence of phages (16). In addition, pre-colonization of fish with phage significantly slowed down the infection and reduced the mortality of rainbow trout (17).

One of the biggest challenges for phage therapy is the imposed selection for phage resistance among phage-exposed bacteria. Bacteria have developed a variety of phage defence strategies, including surface modification and cell aggregation, inactivation of intruding phage DNA by Restriction-Modification and CRISPR-Cas systems, proteolytic digestion of phage particles, and quorum sensing regulation of phage receptor expression (e.g. 18-20). The prevalence and control of these resistance mechanisms depend specifically on the phage-bacterium interaction, on the type and function of the receptor, and the costs of engaging the different mechanisms under various environmental conditions. In many pathogenic bacteria the cell surface molecules are functioning as virulence factors, and phage-driven changes in these structures leading to phage resistance often lead to simultaneous reduction in virulence (21). This trade-off has been detected also among several bacterial fish pathogens, e.g. in *Pseudomonas plecoglossicida* (22), *F. psychrophilum* (23) and *Vibrio anguillarum* (24).

Exposing *F. columnare* to phages has been observed to cause a change in colony morpohotype from the ancestral rhizoid form to rough, which is associated with loss of gliding motility and virulence (25-27). Since a change in colony morphology and loss of virulence have been observed previously also by deletion of genes in the Type IX secretion system involved in gliding motility of *F. columnare* (28), it is likely that mutations in this secretion system are also linked with phage resistance in *F. columnare* (29). Yet, the exact mechanisms by which phages cause the colony morphology change in *F. columnare*, and the functional implications for the bacteria have not been previously explored.

Understanding the mechanisms and consequences of phage resistance in the target bacteria is central for development of successful phage therapy. Thus, in this study, we exposed two *F. columnare* isolates (FCO-F2 and FCO-F9) separately to three different phages, and studied infection dynamics, bacterial viability and colony morphology, and isolated phage-resistant bacteria. Twenty-four phage-exposed and no-phage control isolates were further characterized for their phage resistance, antibiotic susceptibility, motility, adhesion and biofilm formation on polystyrene surface, protease (elastinase, gelatinase and caseinase) activity, virulence on rainbow trout fry, and whole genome sequence. Our results show, that if phage resistance in *F. columnare* is gained via surface modification leading to morphotype change, virulence decreases. However, if the colony morphology remains rhizoid, the isolates remain highly virulent with reduced sensitivity to phage compared to the ancestral wild-type strain.

## Results

### Isolates from phage-exposures: growth, colony morphology and phage resistance

In all phage-exposure cultures of FCO-F2, there was a strong initial phage control of the host population during the first day in all the phage-exposed cultures compared with control culture without phages (Figure 1a). After this, the bacterial density started to recover. The phage-free cultures grew exponentially during the first day, after which they reached a plateau phase. Along with the population decline on day 1, bacterial colony morphotype changed from ancestral rhizoid to rough (Figure 2). From day 1 onwards, more than 88% of the colonies formed by phage-exposed bacterial isolates were rough, the amount reaching at least 97% at the end of the experiment (Figure 1c). In addition, in FCOV-F25 exposure, few soft colonies were observed on day 2 (Figure 2), and in no-phage control cultures, some rough colonies appeared among the prevailing rhizoid ones.

**Figure 1.**
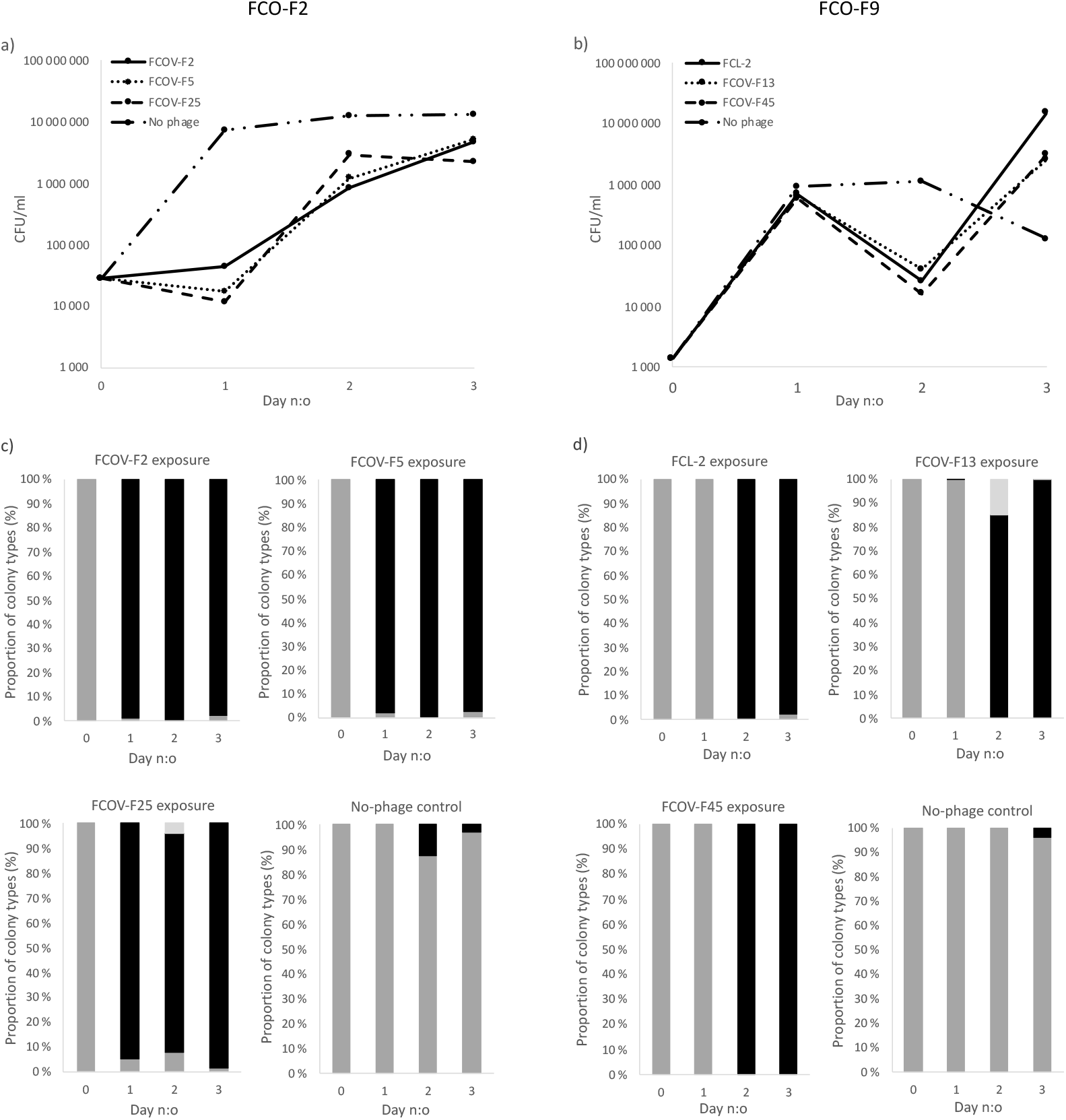
Bacterial growth (a and b), determined as colony forming units mL^−1^, and proportion (%) of different colony types (c and d) of *Flavobacterium columnare* isolates FCO-F2 (a and c) and FCO-F9 (b and d) during the 3-day exposure to phages FCOV-F2, FCOV-F5, FCOV-F25, FCL-2, FCOV-F13 and FCOV-F45. Dark grey bar: proportion of isolates forming rhizoid colony morphology, black bar: proportion of isolates forming rough colony morphology, light grey bar: proportion of isolates forming soft colony morphology.

**Figure 2.**
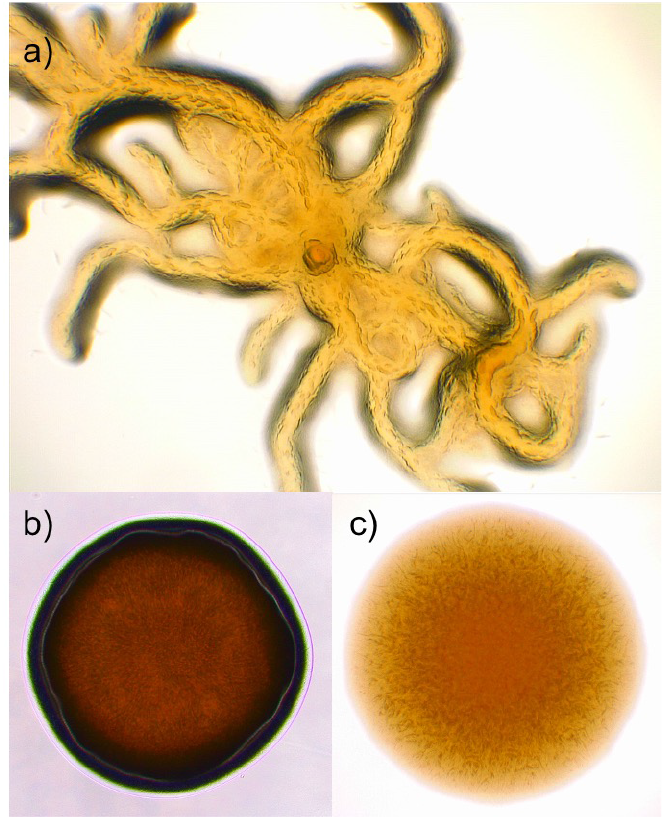
Different colony morphologies formed by *Flavobacterium columnare* on Shieh-agar plates after phage exposure: a) rhizoid, b) rough and c) soft.

FCO-F9 showed slightly different growth dynamics. The bacterial population size increased exponentially during the first day in all cultures (Figure 1b), but decreased drastically on day 2 in response to phage exposure, and then reached exponential growth again. The phage-free cultures reached a plateau phase on day 2, after which the amount of culturable bacteria decreased. From the day 2 population crash and onwards, more than 85% of the colonies formed by phage-exposed bacteria had rough morphology (Figure 1d). At the end of the experiment, more than 98% of the colonies where rough. In FCOV-F13 exposure, a few rough colonies were observed already on day1 and some soft colonies on days 2 and 3. In no-phage control cultures, some (4 %) rough colonies appeared among the rhizoid ones on day 3.

Out of 189 colonies collected from phage exposures, 20 phage-exposed and 4 no-phage control isolates were characterized further (Table 1). Of these isolates, the no-phage control isolates all formed rhizoid colonies similar to their wild-type parent, phage-sensitive isolates FCO-F2 and FCO-F9. Most of the phage-exposed isolates were of rough colony morphology, but F2R58, F2R66 and F9R56 had rhizoid, and F9R69 soft colony morphology.

**Table 1.**
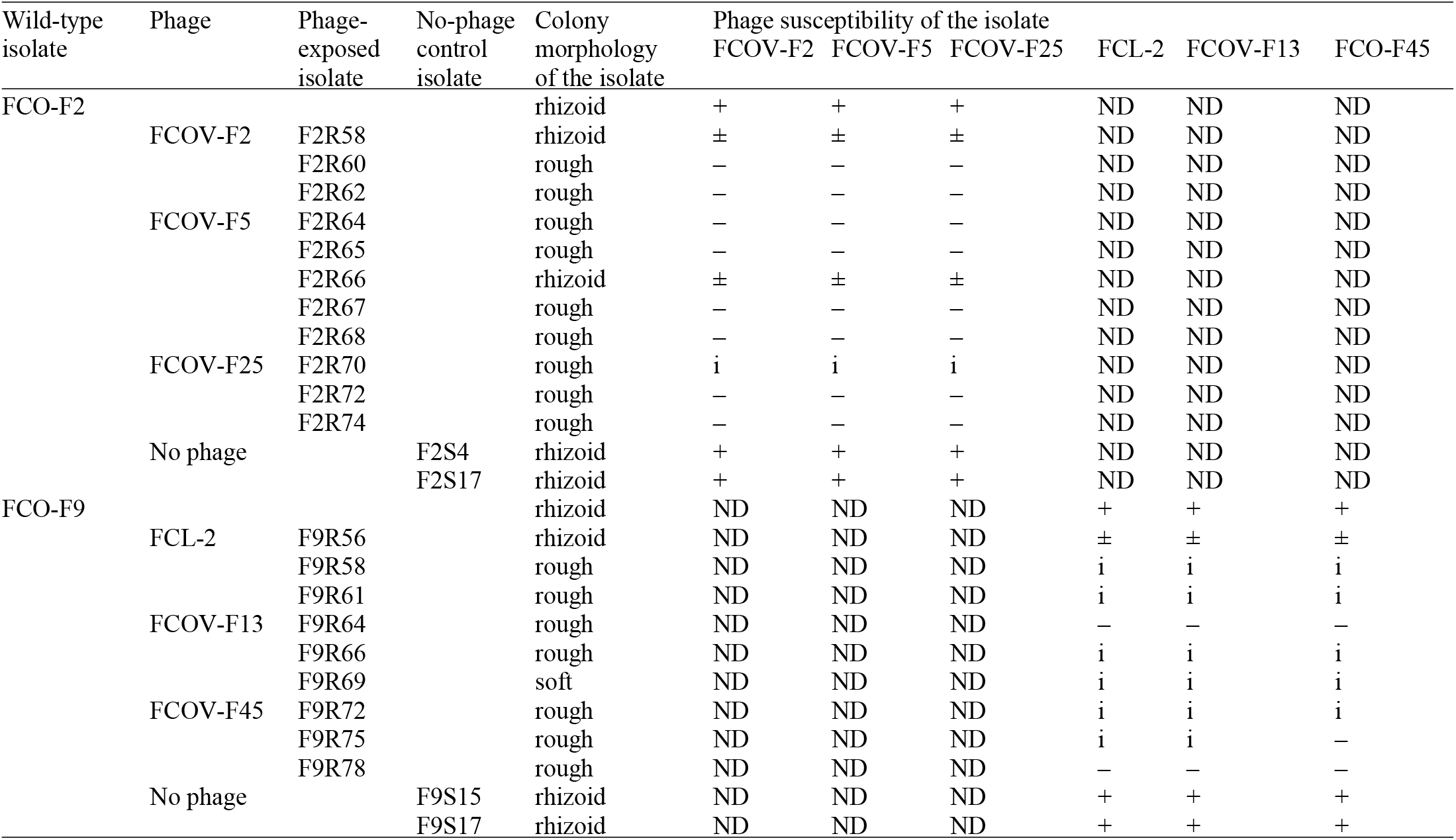
Experimental setup of phage exposure of two phage-sensitive wild-type *Flavobacterium columnare* isolates FCO-F2 (high-virulence, genotype C; exposed for phages FCOV-F2, FCOV-F5 and FCOV-F25) and FCO-F9 (medium-virulence, genotype G; exposed for phages FCL-2, FCOV-F13 and FCOV-F45), and colony morphologies and phage susceptibilities of the 20 phage-exposed (F2R- and F9R-) and 4 no-phage control isolates (F2S- and F9S-) obtained from the exposure cultures. The isolates are shown according to the phage they were exposed to. The susceptibility of the isolates to phages used in exposures: + = sensitive, – = resistant, ± = sensitivity decreased compared to the parent wild-type isolate, i = inhibition of bacterial growth, considered as phage resistance, ND = not determined.

All the phage-exposed rough isolates were resistant to all the phages used to infect the ancestor wild-type bacteria (Table 1). In addition, in some cases, phage caused inhibition of bacterial growth, considered as phage resistance because no clear plaques due to phage infection were detected. The rhizoid phage-exposed isolates turned out to be partly phage-resistant with a 5.5 × 10^5^ to 11 × 10^5^-fold reduction in phage susceptibility compared to the wild type isolates, depending on the specific phage (results not shown). Throughout this paper, these isolates with decreased phage sensitivity are grouped together with the phage-sensitive isolates.

### Antibiotic susceptibility

All isolates showed antibiotic susceptibility patterns similar to the parent wild-type isolates, and no notable differences were observed (Figure S1 and Table S1).

### Motility, adhesion and biofilm formation

Phage-sensitive bacteria forming rhizoid colonies were significantly more motile (determined as colony spreading) than phage-resistant rough or soft morphotypes, irrespective of isolation history (F2-isolates: *P* < 0.001, Oneway ANOVA, LG10 transformation; F9-isolates: *P* ≤ 0.004, Mann-Whitney test) (Figure 3).

**Figure 3.**
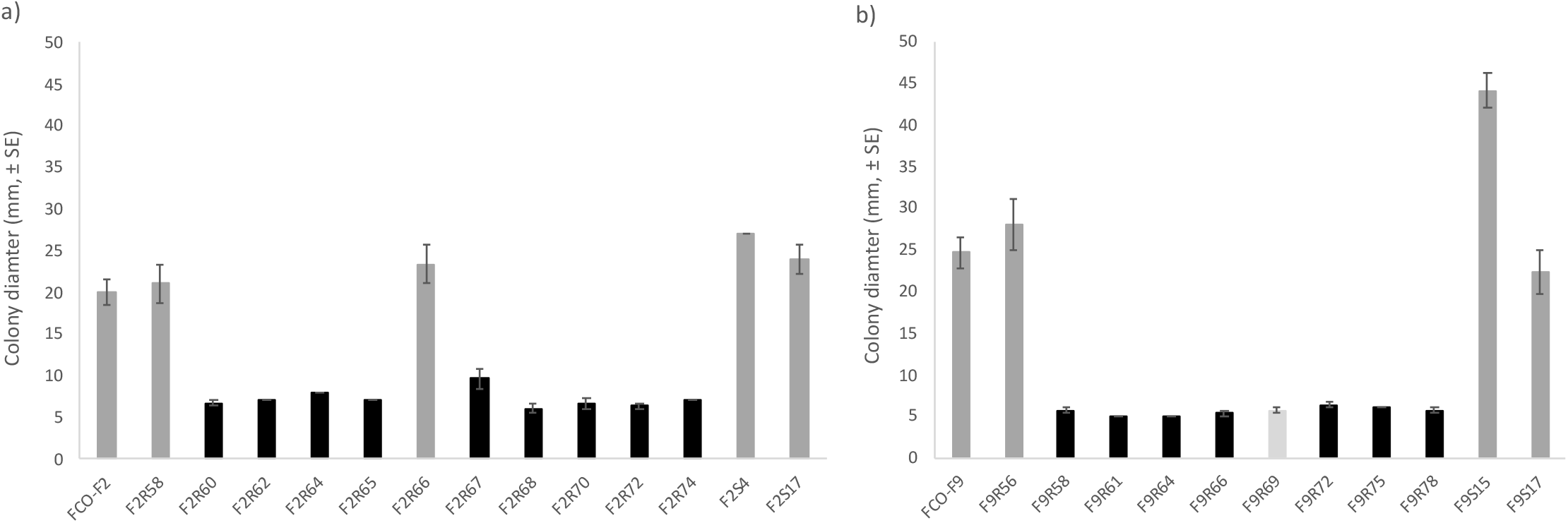
Motility of *Flavobacterium columnare* wild-type a) FCO-F2 and b) FCO-F9 isolates, and their phage-exposed (F2R- and F9R-) and no-phage control (F2S- and F9S-) isolates expressed as colony diameter (mm, ±SE) on TYES agar. All the phage-sensitive rhizoid colonies forming isolates (dark grey bar) were significantly more motile than phage-resistant rough (black bar) or soft (light grey bar) morphology isolates (F2-isolates: P < 0.001, Oneway ANOVA, LG10 transformation; F9-isolates: P ≤ 0.004, Mann-Whitney test).

Compared to the parent wild-type FCO-F2 isolate, there was a large variability on the adhesion capacity of individual phage-resistant F2-isolates (Figure 4a). Phage susceptibility (rhizoid vs. rough colony type) or phage used in the co-culture experiment did not influence bacterial adhesion capacity (*P* = 0.3: Mann-Whitney test and *P* = 0.564: Kruskal-Wallis test, respectively).

**Figure 4.**
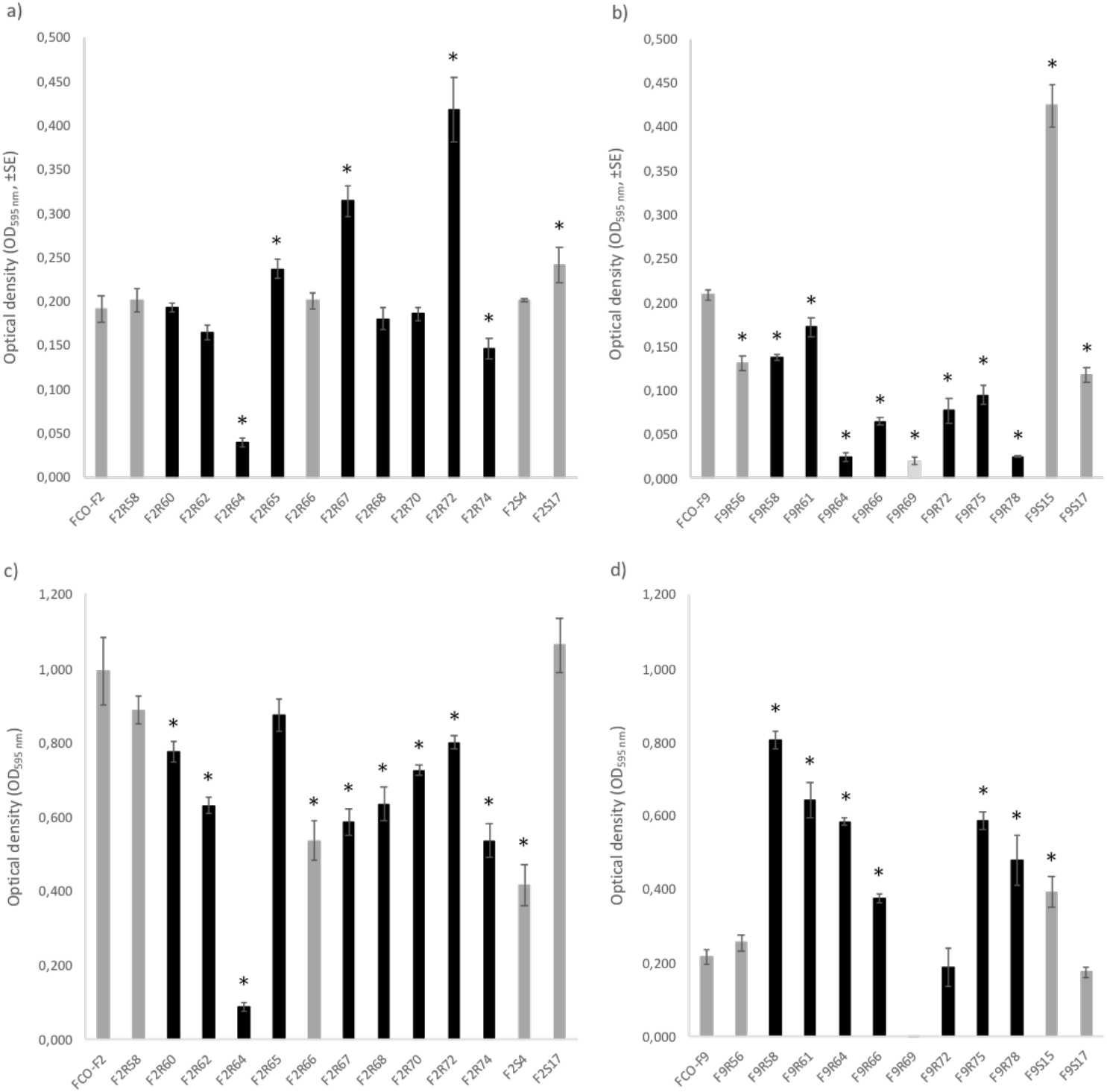
Adherence (a and b) and biofilm forming (c and d) capacity of *Flavobacterium columnare* wild-type FCO-F2 (a and c) and FCO-F9 (b and d) isolates, and their phage-exposed (F2R- and F9R-) and no-phage control (F2S- and F9S-) isolates on a polystyrene surface measured as optical density (OD_595 nm_, ±SE). Asterisks indicate the statistically significant difference (P < 0.05) compared to the parent wild-type isolate. F9S69 did not form any biofilm and was thus excluded from the statistical analyses. Dark grey bars: phage-sensitive isolates forming rhizoid colony morphology, black bars: phage-resistant isolates forming rough morphology, light grey bar: phage-resistant isolate forming soft morphology.

Most of the individual phage-exposed and no-phage control F2-isolates had significantly lower biofilm forming capacity than in parent wild-type FCO-F2 (*P* ≤ 0.017: Oneway ANOVA, LDS multiple comparisons, square root transformation) (Figure 4c). Still, there was no statistical difference in biofilm formation between phage-sensitive rhizoid and resistant rough morphology F2-isolates (*P* = 0.062: Oneway ANOVA).

Again, the bacterial strain F9 behaved differently compared to F2. In contrast to the phage-resistant F2-isolates, the phage-resistant rough and soft morphology F9-isolates had significantly lower adherence than sensitive rhizoid isolates (*P* < 0.001: Oneway ANOVA, LDS multiple comparisons, square root transformation) (Figure 4b). In addition, isolates exposed to phages isolated in 2017, FCOV-F13 and FCOV-F45, had significantly lower adhesion capacity than in isolates exposed to FCL-2 isolated in 2009 (*P* < 0.001: Mann-Whitney test). This may indicate phage FCL-2 uses different phage receptor (see later).

In contrast to adhesion ability, biofilm forming capacity of the most of the individual phage-exposed and no-phage control F9-isolates was significantly higher compared to wild-type parent isolate (*P* ≤ 0.004: Oneway ANOVA, LDS multiple comparisons) (Figure 4d). F9R69 with soft colony morphology did not form any biofilm and thus excluded from the multiple comparisons. Phage-resistant rough F9-isolates had significantly higher biofilm forming capacity than sensitive rhizoid morphotypes (*P* < 0.001: Oneway ANOVA, square root transformation).

### Protease activity: elastinase, gelatinase and caseinase

Elastinase activity was detected in the wild-type, and all the phage-sensitive rhizoid FCO-F2 isolates and one resistant rough F2-isolate (clear zone ratio > 1), whereas all remaining resistant, rough morphology isolates, had completely lost the ability to degrade elastin (Figure 5a). There were no differences in elastinase activity between the elastinase positive isolates (*P* = 0.843: Oneway ANOVA). Elastinase activity was not detected in any of the F9-isolates (clear zone ratio = 1) (Figure 5b).

**Figure 5.**
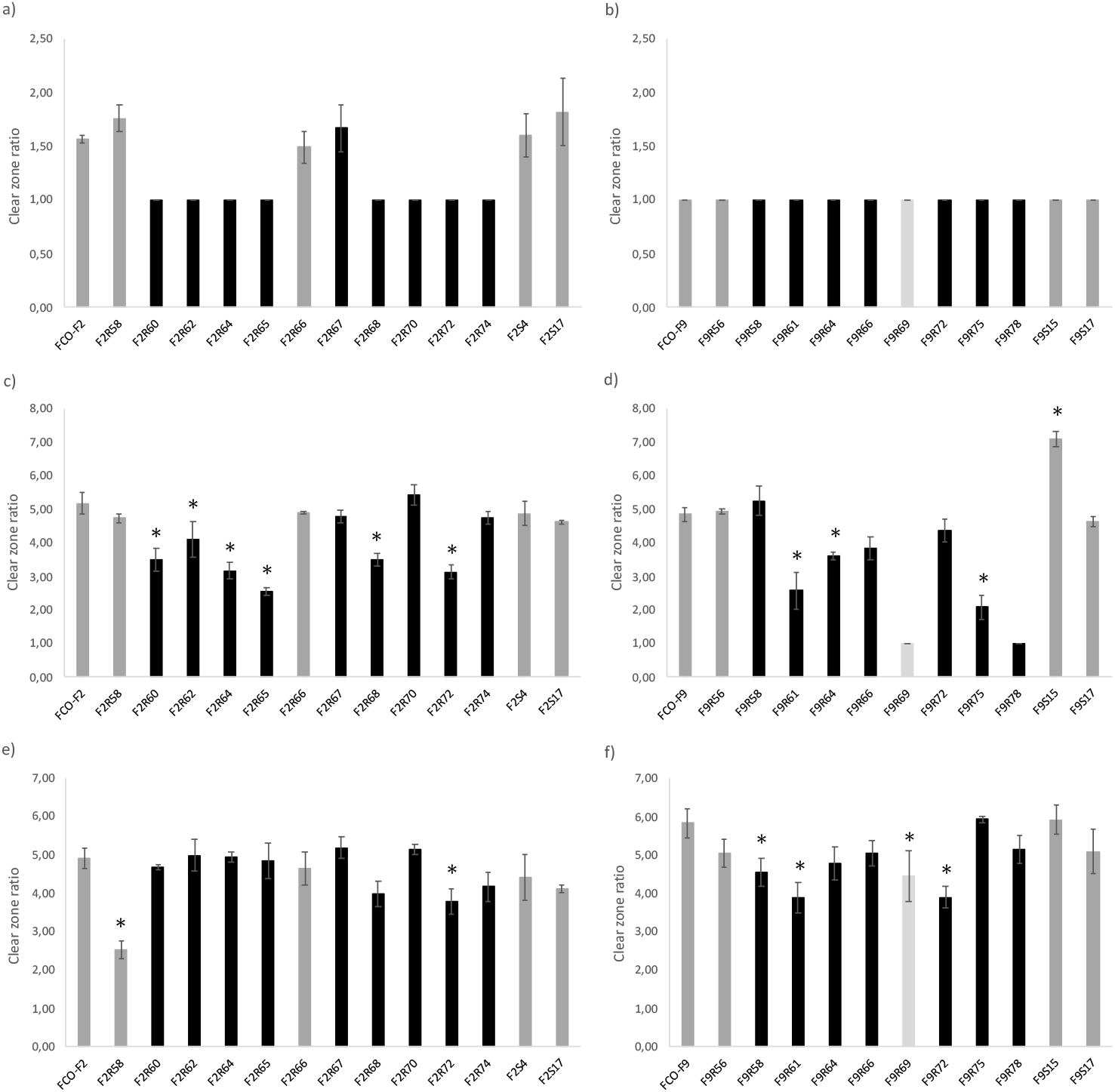
Protease (elastinase: a and b, gelatinase: c and d, and caseinase e and f) activity of the *Flavobacterium columnare* FCO-F2 (a, c and e) and FCO-F9 (b, d and f) isolates, and their phage-exposed (F2R- and F9R-) and no-phage control (F2S- and F9S-) isolates. The activity was measured as the clear zone ratio (clear zone diameter/colony diameter, ±SE) on TYES agar supplemented with elastin, gelatin and skim milk (caseinase). The asterisk indicates significant reduction in protease activity (P < 0.05) compared to the parent wild-type isolate. A clear zone ratio 1 indicates no protease activity. Isolates with no activity were excluded from the statistical analyses. Dark grey bars: phage-sensitive isolates forming rhizoid colony morphology, black bars: phage-resistant isolates forming rough morphology, light grey bar: phage-resistant isolate forming soft morphology.

There were variations in gelatinase activity between individual F2- and F9-isolates (Oneway ANOVA, LDS multiple comparisons) (Figure 5c and d). However, among both F2- and F9-isolates, gelatinase activity of phage-resistant rough morphotypes was lower than that of sensitive rhizoid morphotypes (F2-isolates: *P* = 0.018, Oneway ANOVA, exponential transformation; F9-isolates: *P* < 0.001, Oneway ANOVA). Two of the phage-exposed F9-isolates (F9R69 and F9R78) did not have any gelatinase activity and were thus excluded from the multiple comparisons

Less variation in caseinase activity between individual isolates was observed (Oneway ANOVA, LDS multiple comparisons) (Figure 5e and f), and phage-sensitive rhizoid and resistant rough F2-isolates did not differ from each other (*P* = 0.058: Oneway ANOVA. On the other hand, caseinase activity of phage-resistant rough and soft F9-isolates was lower than that of sensitive rhizoid isolates (*P* = 0.007: Oneway ANOVA).

### Virulence

Rainbow trout fry were exposed to wild-type, phage-exposed and no-phage control isolates, and all of them caused mortality during 24 h (Figure 6). The phage-sensitive rhizoid morphotypes were most virulent, causing 100 % mortality, whereas resistant rough and soft morphotypes were less virulent, causing 46.7 % mortality at highest (except for phage-resistant rough morphotype F2R70, which caused 100 % mortality). Mortality of control fish was 15 %, but no bacterial growth was observed from these fish. However, *F. columnare* growth was observed from all the fish exposed to bacteria. Colony morphotype of the bacterial isolates did not change during the infection.

**Figure 6.**
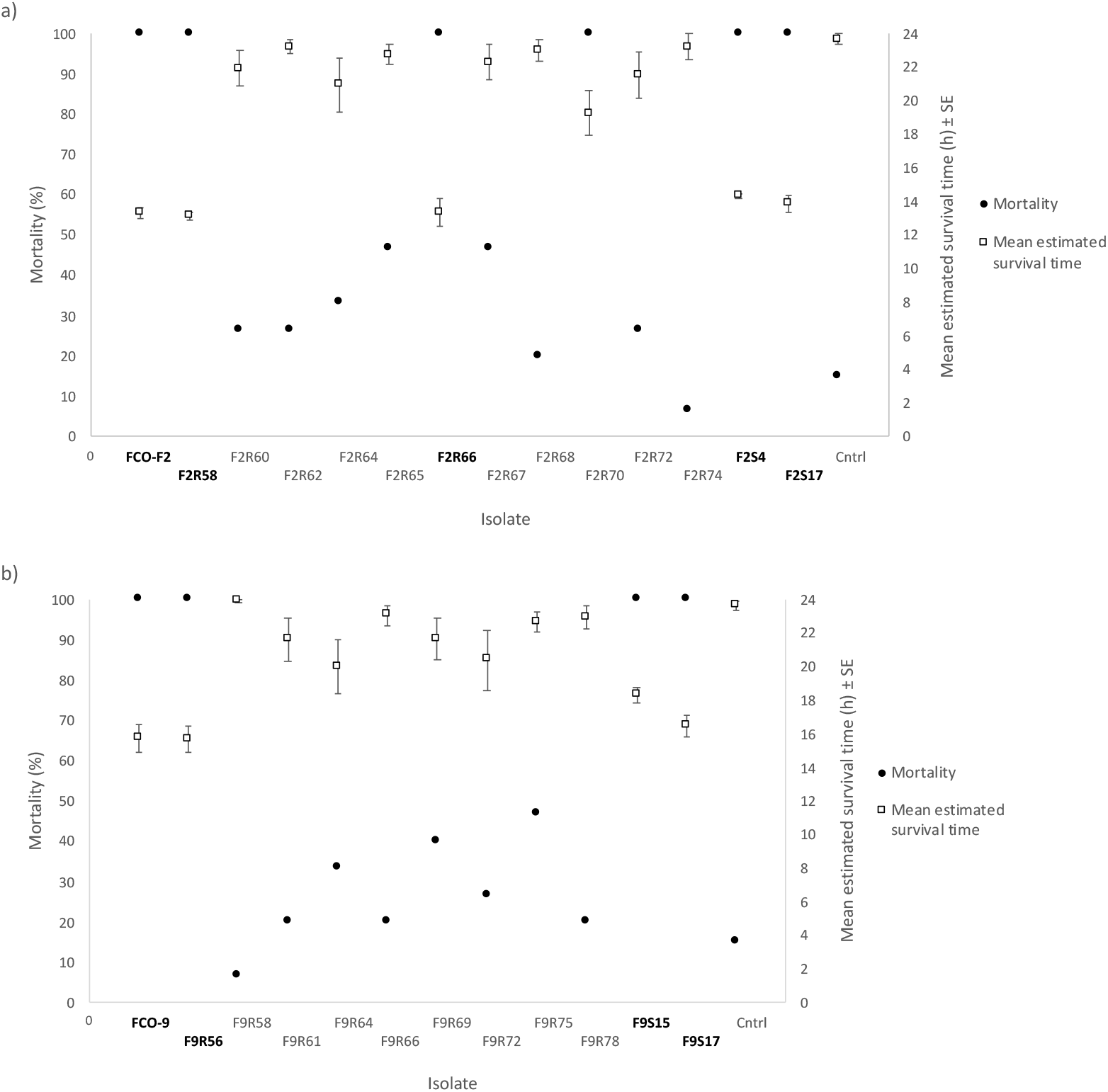
Mortality percent and estimated survival time (± SE) of rainbow trout (*Oncorhynchus mykiss*) during 24-h experimental infection with wild-type *Flavobacterium columnare* FCO-F2 (a) and FCO-F9 (b), and their phage-exposed (F2R- and F9R-) and no-phage control (F2S- and F9S-) isolates. Phage-sensitive rhizoid colonies forming isolates are written bold. Cntrl = control with no bacterial infection.

When comparing the data according to the phage susceptibility and thus colony morphology, cumulative mortality of fish infected with phage-sensitive rhizoid morphotypes, irrespective of if they were wild-type, phage-exposed or no-phage control isolates, was significantly higher than mortality caused by phage-resistant rough or soft morphotypes among both F2 and F9 isolates (*P* < 0.001, Kaplan-Meier Survival Analysis). Also, the estimated survival time (Kaplan-Meier Survival Analysis) was shortest in fish infected with sensitive rhizoid isolates (Figure 6). In case of F2-isolates, mortality caused by phage-resistant rough isolates was also significantly higher than mortality of control fish, but mortality caused by resistant rough and soft F9-isolates did not differ from each other or from the control fish mortality. Mortality caused by rhizoid phage-sensitive F2 isolates started to peak at 12 hours post infection (p.i.) and in F9 at 16 hours p.i. (*P* < 0.001, Kaplan-Meier Survival Analysis), but between rough phage-resistant F2 and F9 isolates the mortality patterns were more similar starting to increase slowly at 2-3 hours p.i. (*P* = 0.217, Kaplan-Meier Survival Analysis). However, there were differences in cumulative mortalities caused by individual isolates in each morphology group (Data set S1).

### Whole genome sequencing

Genome data of wild-type *F. columnare* isolates FCO-F2 and FCO-F9 is presented in Table 2.

**Table 2.** Data on genomes of wild-type *Flavobacterium columnare* strains FCO-F2 and FCO-F9.

| Wild-type isolate | Genetic group | Genome size (bases) | N:o of ORFs | GC % |
| --- | --- | --- | --- | --- |
| FCO-F2 | C | 3 221 312 | 3 280 | 31.7 |
| FCO-F9 | G | 3 261 403 | 3 374 | 31.7 |

Genomic comparisons between F2 wild type and phage-exposed isolates revealed a limited number of genomic changes. In seven out of 11 isolates, single mutation leading to formation of wrong or truncated proteins was observed in the phage-resistant mutants (Table 3).

**Table 3.** Mutations revealed by whole genome sequencing (Illumina) in F2 phage-exposed *Flavobacterium columnare* isolates compared to their wild type (wt) isolate FCO-F2. The isolates are shown according to the phage they were exposed to. CDS = coding sequence, → = change to, Del = deletion, Ins = insertion, aa = amino acid.

| (Phage) | Phage-exposed isolate | Colony morphology | Gene/CDS | Mutation | Location (base n:o) in wt genome | Outcome |
| --- | --- | --- | --- | --- | --- | --- |
| (FCOV-F2) | F2R58 | rhizoid | rlmF | T → A | 21 350 | No aa change |
|  | F2R60 | rough | sprA | Ins GT | 1 314 323 – 1 314 324 | Change in reading frame → stop codon<br>→ two truncated proteins |
|  | F2R62 | rough |  |  |  |  |
| (FCOV-F5) | F2R64 | rough | sprA | Ins G | 1 317 523 | Change in reading frame → stop codon<br>→ two truncated proteins |
|  | F2R65 | rough | sprA | Ins G | 1 317 524 | Change in reading frame → stop codon<br>→ two truncated proteins |
|  | F2R66 | rhizoid |  |  |  |  |
|  | F2R67 | rough | gldB | Del T | 1 122 801 | Truncated/wrong protein |
|  | F2R68 | rough |  |  |  |  |
| (FCOV-F25) | F2R70 | rough | OmpH family outer membrane protein | Ins G | 1 275 242 | Change in reading frame<br>→ wrong protein |
|  | F2R72 | rough | gldN | Ins TCTAC | 1 013 274 – 1 013 278 | Change in reading frame → stop codon<br>→ two truncated proteins |
|  | F2R74 | rough | sprA | Del A | 1 313 911 | Change in reading frame → stop codon<br>→ two truncated proteins |

Notably, the majority of the mutations were located in genes coding for gliding motility proteins *gldB* (F2R67), *gldN* (F2R72) and *sprA* (F2R60, F2R64, F2R65, F2R74). Isolate F2R70 had one nucleotide insertion in OmpH family outer membrane protein coding gene. Three isolates (F2R62, F2R66, F2R68) did not show any genomic changes relative to the wild type. In isolate F2R58 with decreased phage sensitivity, one nucleotide change in *rlmF* gene (coding for rRNA large subunit methyltransferase F) did not lead to amino acid change. No mutations were observed in the no-phage control isolates. At certain points of ribosomal RNA operons in all phage-exposed and no-phage control isolates, and also in a 736 221 bp sequence (hypothetical protein coding sequence in wild-type FCO-F2 genome used as a reference) in phage-exposed isolates F2R66 and F2R68, there was a poor coverage of reads leading to unclear sequences, which prevented detection of possible mutations in this region.

In F9 phage-exposed isolates, one or two mutations per isolate in all the other isolates, except for F9R58, were observed (Table 4). Mutations in isolates exposed to FCOV-F45 had insertions whereas FCOV-F13 exposed isolates had deletions or single nucleotide chances in genes coding for gliding motility proteins *gldG* (F9R72), *gldM* (F9R64, F9R69, F9R78) and *gldN* (F9R69, F9R75), leading to formation of wrong or truncated proteins. Interestingly, in the isolate F9R69 (exposed to FCOV-F13) with a soft colony type, a deletion of genomic region of 4 701 bp was observed, spanning over gliding motility genes *gldM* and *gldN*, and sequences coding for FAD-binding oxidoreductase, DUF3492 domain-containing protein and a hypothetical protein (Figure 7).

**Table 4.**
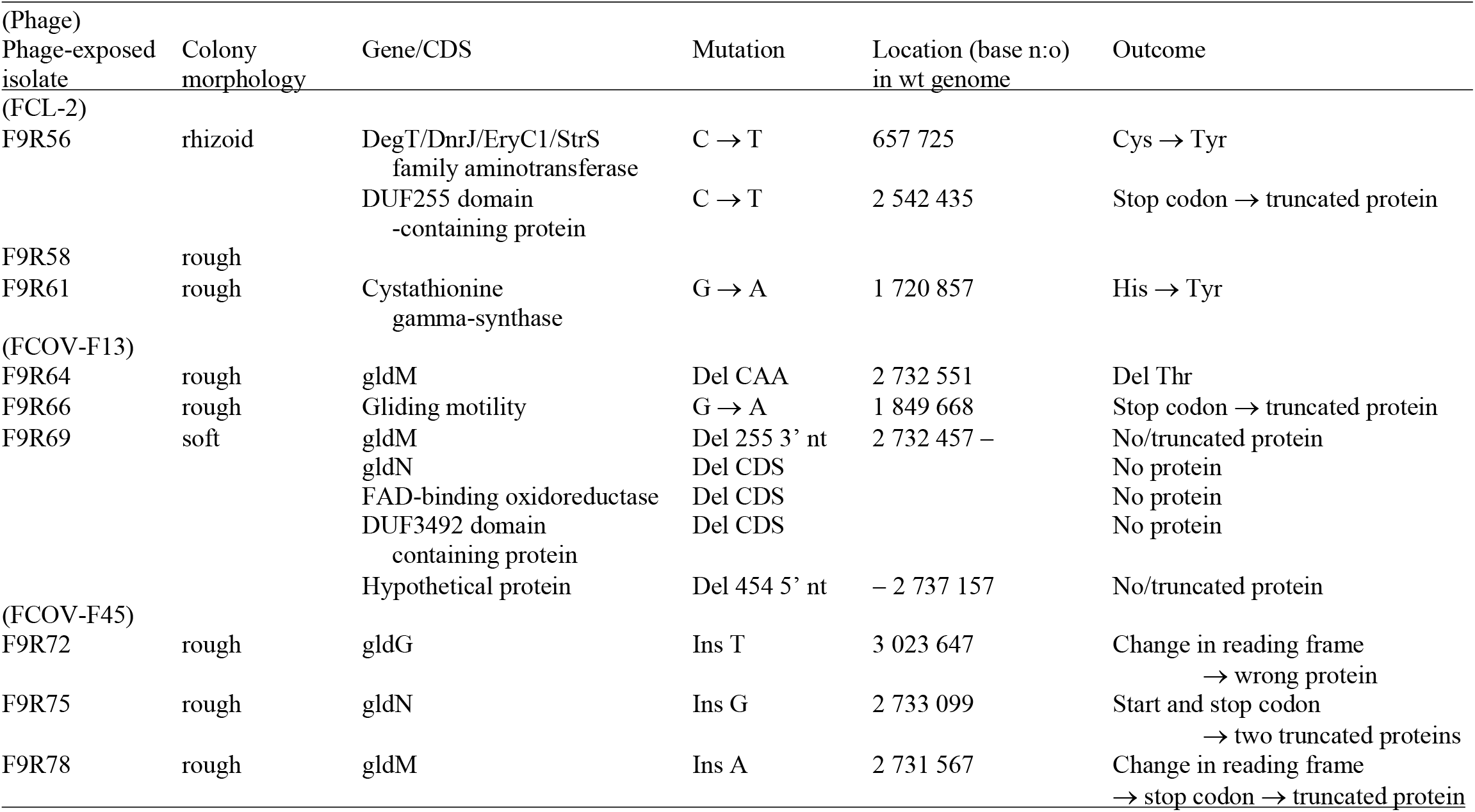
Mutations revealed by whole genome sequencing (Illumina) in F9 phage-exposed *Flavobacterium columnare* isolates compared to their wild type (wt) isolate FCO-F9. The isolates are shown according to the phage they were exposed to. CDS = coding sequence, → = change to, Del = deletion, nt = nucleotide, Ins = insertion, aa = amino acid

**Figure 7.**
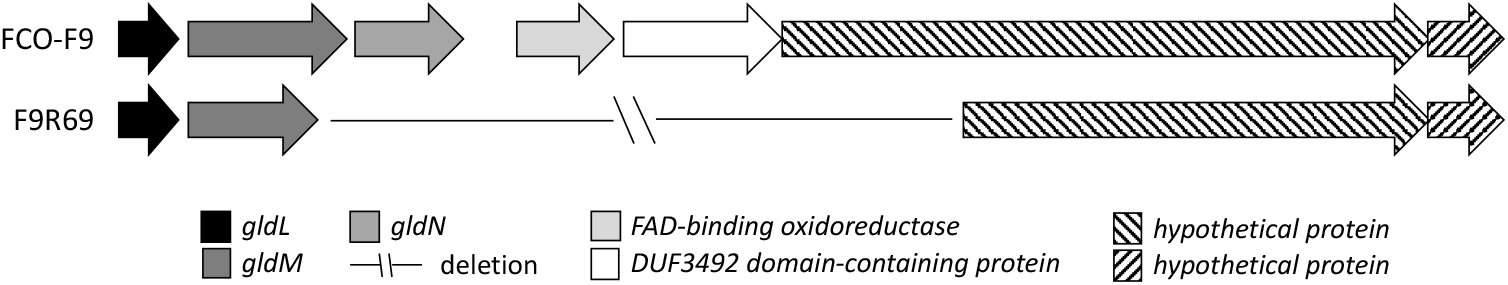
Deletion of genomic region covering 4 701 bp in FCOV-F13 exposed, soft colonies forming, phage-resistant *Flavobacterium columnare* isolate F9R69.

On the contrary, no mutations in gliding motility genes were observed in F9 isolates exposed to FCL-2, but instead, two of these isolates had one nucleotide change in DegT/DnrJ/EryC1/StrS family aminotransferase and DUF255-domain containing protein (F9R56), and cystathionine gamma-synthase (F9R61) coding genes, leading to either one amino acid change or truncated protein. No mutations were observed in no-phage control isolates. Around 2 000 620 bp (hypothetical protein coding sequence in B185 genome used as a reference), there was a poor coverage of reads leading to unclear sequence in both wild type FCO-F9, phage-exposed and no-phage control isolates, which prevented detection of possible mutations in this region.

## Discussion

Phage therapy is seen as an attractive option to treat and prevent bacterial diseases, but the development of phage resistance in target bacteria is considered as one of the main problems related to the use of phages. Our results describe the selection for phage resistance in two different *F. columnare* isolates upon exposure to six specific phages. We show that phage resistance is associated with reduction in virulence and virulence-related phenotypic changes in the bacterium. Our genetic data indicate that in most cases phage resistance is caused by surface modifications, often related to the type IX secretion system connected to flavobacterial gliding motility machinery. Mutations in the genes coding for an outer membrane protein or genes related to gliding motility seem to be phage specific and likely prevent phage attachment, possibly in a phage specific manner, and lead to morphology change and loss of virulence.

In the present study, phage-exposure caused significant changes in bacterial phenotypic characteristics (motility, adhesion, protein secretion and virulence - details below) leading to phage resistance. In most isolates, these changes could be linked to changes in gliding motility-related genes. Flavobacteria show gliding motility on surfaces (29), and mutations in any of the genes coding for gliding motility machinery proteins have been shown to lead to loss of motility (e.g. 30, 31). Gliding is also connected to virulence, since part of the gliding motility machinery (GldK, GldL, GldM, GldN, PorV, SprA, Spr E, SprF and SprT) is used as a type IX secretion system found in Bacteroidetes (28, 32). Indeed, phage resistance due to loss of motility has been linked with decreased virulence in *F. columnare* also previously (27), and *F. columnare gldN* mutants have been shown to exhibit both decreased proteolytic and chondroitinase activity, and virulence on rainbow trout (28). Similarly, phage resistance was associated with loss of motility and mutations in genes related to cell surface properties and gliding motility in *F. psychrophilum* (23) and in *F. johnsoniae* (31, 33). Together, the results suggest that the type IX secretion system is a key target for infection by a wide range of phages and across the Flavobacterium genus, and that mutations leading to morphology changes and loss of motility is a general response to phage exposure in this bacterial group.

Exposure to a specific phage led to different mutations in gliding motility genes in different *F. columnare* isolates, as also seen in phage-resistant *F. psychrophilum* (23), indicating that several genes are involved in phage attachment and infection of *F. columnare* phages. Furthermore, genomic analysis of one soft colony isolate revealed a large deletion (4 701 bp), spanning over two gliding motility genes. However, although all rough colony forming isolates were phage-resistant, not all these isolates (F2R62, F2R66, F2R68 and F9R58) had mutations in genes coding for proteins related to gliding motility, or elsewhere in their genome. This may indicate that development of phage resistance and colony morphology change are also influenced by gene expression or epigenetic modifications, leading to variation in colony morphology, as suggested previously (34). For example, in *Bordetella* spp, phage resistance is regulated via phase variation in virulence related factors, such as some adhesins, toxins and type III secretion system (reviewed in 35). Interestingly, isolates exposed to FCL-2 did not have mutations in gliding motility related genes, suggesting that FCL-2 uses other receptors for infection of *F. columnare* than the other phages. FCL-2 differs genetically from other phages infecting genetic group G bacteria (This article was submitted to an online preprint archive [36]), supporting this suggestion.

Generally, point mutations and changes in receptor expression enable a rapid and efficient response of bacterial populations to phage exposure. However, the large phenotypic costs of mutational derived phage resistance observed in *F. columnare* in this study suggest that these mutations may be dynamic and most probably also rapidly reverting back to the sensitive form in *F. columnare*. Indeed, reversion of both phage-driven and spontaneously formed rough colony types back to rhizoid has been observed to happen in *F. columnare* subcultures (27). Various mechanisms to regain phage resistance have been found also in fish pathogenic *F. psychrophilum* (23) and *V. anguillarum* (24), in which a rapid reversion back to phage-sensitive phenotype has been shown to occur. This sort of dynamics in phage resistance has also been observed in a human symbiont *Bacteroides thetaiotaomicron* (37), suggesting that the phenomenon may be common among wide variety of bacteria.

Phage-exposed *F. columnare* isolates F2R56, F2R66 and F9R58 did not respond to phage infection with surface modifications, but maintained their original rhizoid colony morphotype and high virulence. These rhizoid isolates were not completely resistant to phage, although phage infection efficiency dropped markedly (up to a million-fold decrease), suggesting some other mechanism for reducing infection efficiency. *F. columnare* has two functional CRISPR systems, which have been shown to adapt under phage exposure at fish farms (38). However, we did not observe additional CRISPR spacers in whole genome sequencing. The same was observed in phage exposed *F. psychrophilum* isolates in which no differences to the wild-type strain’s CRISPR composition were found (23). In our experience, CRISPR adaptation in *F. columnare* requires different experimental set-up with longer co-culture time in low nutrient medium, followed by enrichment in high-nutrient medium (This article was submitted to an online preprint archive [39]). Thus, the decreased phage sensitivity of rhizoid phage exposed isolates most probably is a consequence of yet unknown functions which need to be studied in the future.

In addition to type IX secretion system, also type I and VI secretion systems are known to function in *F. columnare* (40). Possible secretion of virulence related factors through type I and VI secretion systems in *F. columnare* could be one of the reasons why also rough phage-resistant isolates caused some mortality in fish, and explain their gelatinase and caseinase activity despite morphology change. It has also been shown recently, that virulence of *F. columnare* increases in the mucus and with increasing mucin concentration (17). As the mucus-covered fish surface is the main infection route of *F. columnare*, it is probable that some *F. columnare* virulence factors, such as proteinase activity, are expressed differently in growth media compared to the *in vivo* infection situation. This possible differential expression could also explain the mortality caused by phage-resistant rough isolates.

The ability to adhere and form biofilm has a major role in bacterial infections and in colonizing niches (41). In *F. columnare*, adhesion and biofilm forming capacity may have a central role in their persistence in the farming environment (e.g. tanks and water systems) (42), but also in establishing the first steps of infection on the fish surfaces (43). Our results indicate that *F. columnare* strains differ in their adherence and biofilm forming characteristics. Whereas phage exposure had no clear effect on the adhesion capacity of the F2-isolates, phage resistance led to decrease in biofilm forming capacity in most of the individual phage-resistant F2-isolates. This is in agreement with the systematic reduction in biofilm forming properties of phage-resistant *F. psychrophilum* relative to the wild type (23). Adhesion capacity of F9 phage-resistant isolates, on the other hand, was significantly lower compared to the wild-type parent isolate, but rough phage-resistant F9-isolates had significantly higher biofilm forming capacity compared to rhizoid sensitive isolates. These results partly differ from what we have found earlier (25, 26), most likely because in the previous studies the rough colonies were formed spontaneously, without phage exposure. Indeed, morphology of spontaneously formed rough colonies and these morphotypes’ ability to move when cultured in low-nutrient media differ from rough morphotypes formed under phage exposure (27). However, together our results indicate, that since *F. columnare* phages are genetically group-specific, they might be using different receptors, which, in turn, causes differences in bacterial resistance mechanisms between genetic groups.

*F. columnare* infections are routinely treated by antibiotics at fish farms. In this study, phage resistance did not affect the antibiotic susceptibility of any of the isolates studied. Lack of association between development of antibiotic resistance and bacteriophage resistance has also been shown e.g. in *Escherichia coli* (44). Based on our results, phage resistance does not increase a risk of antibiotic resistance development, and thus, phage-therapy given as a cure or prophylactic treatment at fish farms most probably does not rule out the possible concomitant use of antibiotics as therapeutic agents against columnaris infections. Indeed, it was shown by using *P. fluorescence* as a model bacterium, that applying phages together with antibiotic treatments may inhibit the evolution of antibiotic resistance in pathogenic bacteria (45).

To summarize, our results show, that even though *F. columnare* rapidly develops phage resistance under phage exposure, the arise of phage resistance does not pose a high risk for a development of phage therapy against columnaris infections in rainbow trout. This is because phage resistance leads to decrease in bacterial virulence, adherence to surfaces and protease secretion. Based on our results with experiments with two genetically different wild-type bacterial isolates, development and regulation of phage resistance in *F. columnare* is a multifactorial process, partly affected by formation of mutations mainly in gliding motility and type IX secretion system related genes, and partly by other defence mechanisms against phages, the function of which needs to be studied in the future.

## Materials and methods

### Bacterial and phage isolates

Bacteria and phages used in this study were isolated from water samples collected from fish farms during columnaris outbreaks (This article was submitted to an online preprint archive [36]) (Table 5). Bacteria were confirmed as *F. columnare* by restriction fragment length polymorphism (RFLP) analysis of 16S rRNA gene and classified into genetic groups by RFLP of 16S-23S internal transcribed spacer (ITS) region (This article was submitted to an online preprint archive [36]). All the six phages belong to the *Myoviridae* family and have been characterised with respect to host range and genomic composition (This article was submitted to an online preprint archive [36]).

**Table 5.**
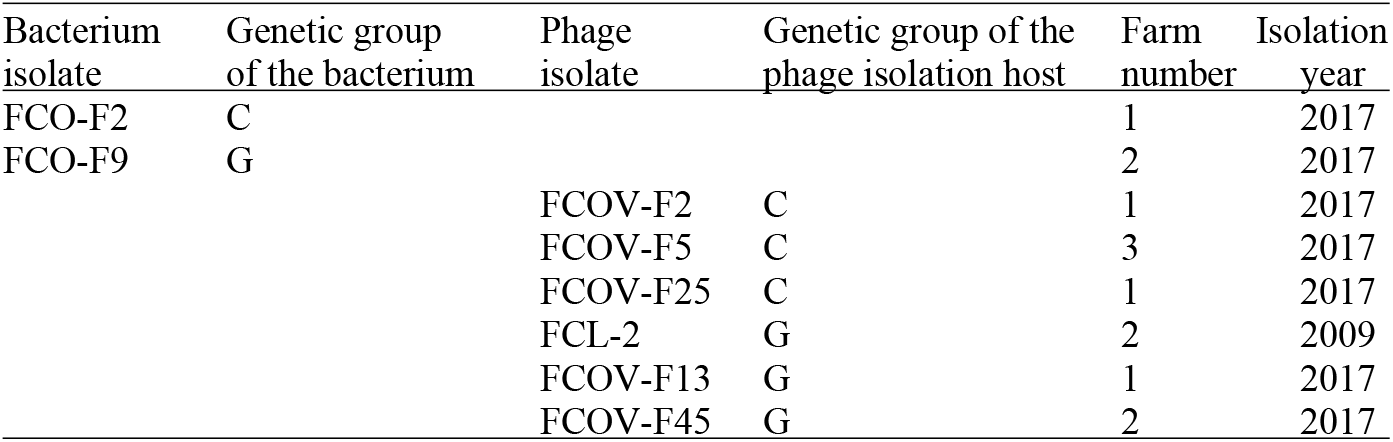
*Flavobacterium columnare* isolates and phages used in this study. Bacteria and phages were isolated from Finnish fish farms. *F. columnare* isolates have previously been categorized into genetic groups by restriction fragment length polymorphism analysis of internal transcribed spacer region between 16S and 23S rRNA genes (This article was submitted to an online preprint archive [36]).

### Bacterial cultures and phage lysates

For phage exposure and virulence test, *F. columnare* isolates were inoculated from cryopreserved (–80°C) stocks in modified Shieh-medium (46) and grown for 48 h at 25°C with 120 rpm agitation. After this, subcultures were made in modified Shieh-medium and grown for 24 h at 25°C with 120 rpm agitation. The optical density (OD) of the bacterial broth suspensions was measured spectrophotometrically at 595 nm and adjusted to 5 × 10^5^ colony forming units (CFU) mL^-1^ for phage exposures and 5 × 10^6^ CFU mL^-1^ for virulence experiment (based on previously determined OD/CFU relationship). For other test, *F. columnare* isolates were cultured in TYES broth (47), washed in TYES broth by centrifugation at 5310 × *g* for 15 min at 4°C, and cultures spectrophotometrically adjusted to OD 0.6 at 520 nm (approximately 10^8^ CFU mL^-1^).

Phage lysates were produced using “double layer agar” -method (48) as follows: Three mL of melted (47°C) top agar (0.5%) including 300 μL of 24-hour subculture of the host bacterium and 100 μL of phage (tenfold dilutions in Shieh medium) was poured on Shieh agar and grown for 48 h at 25 °C. Five mL of Shieh-medium was added on top of Shieh agar plates with confluent lysis and incubated at 7°C for 12-18 h in constant agitation (90 rpm). The lysates were collected, filtered (PES membrane, pore size 0.45 µm, Nalgene®), and stored at +7°C or at –80°C with 20 % glycerol. For phage exposure, phage lysates were diluted with Shieh medium to 5 × 10^5^ plaque forming units (PFU) mL^-1^.

### Phage exposure experiments and isolation of colonies

Two phage-sensitive wild-type *F. columnare* isolates, the high-virulence FCO-F2 isolate (genetic group C) and the medium-virulence FCO-F9 isolate (genetic group G) (This article was submitted to an online preprint archive [36]), were each exposed to three phages in separate experiments with individual phages. Isolate FCO-F2 was exposed to phages FCOV-F2, FCOV-F5 and FCOV-F25, and isolate FCO-F9 to phages FCL-2, FCOV-F13 and FCOV-F45, in accordance with the host range of the phages. Cultures with only bacteria served as no-phage controls. The exposures were carried out in 20 mL of autoclaved fresh water (Lake Jyväsjärvi) in triplicate cultures under constant agitation (120 rpm) at 25°C for three days at a multiplicity of infection (MOI) at inoculation of 1 (1 × 10^4^ CFU and PFU mL^-1^). The cultures were sampled every 24h for three days, by making a serial tenfold dilution of samples, and spreading on Shieh-agar plates. After up to 4 days of incubation at room temperature, CFUs and colony morphologies were determined from the plate cultures. Two to three colonies from each triplicate culture at each sampling point were picked, and pure-cultured directly on Shieh agar plates three times to get rid of any phage contamination. Colonies were then checked for phage resistance by spot assay on agar plates: bacterial laws on top agar were prepared as above and 10 μL of ten-fold diluted original phage lysates (used in initial exposures) were spotted on agar. After 48-h incubation at 25 °C, bacterial plates with no observed plaques or confluent lysis were considered as phage-resistant. Altogether 189 colonies from phage-exposed and no-phage control exposures were isolated from plate cultures. From this collection, 20 phage-exposed and 4 no-phage control isolates were selected for further analysis (Table 1).

The phage-exposed and no-phage control isolates were named according to the latter part of the wild-type bacterial host, a letter R for phage-exposed and S for no-phage control isolate, plus a running number for the isolated colony. For example, F2R2 is the second selected phage-exposed colony of the *F. columnare* wild-type isolate FCO-F2. Correspondingly, the second *F. columnare* isolate from no-phage control cultures was marked as F2S2. For simplicity, wild-type FCO-F2 and all its subsequent isolates from the phage and control exposures are commonly called F2-isolates in this paper. Correspondingly, wild-type FCO-F9 and its subsequent isolates are called F9-isolates.

### Antibiotic sensitivity

Changes in susceptibility of phage-exposed *F. columnare* isolates towards antibiotics was tested using the Kirby-Bauer disc diffusion method (49) on diluted Mueller-Hinton (50) agar medium supplemented with 5 % w/v fetal calf serum. A 40 µL volume of each isolate suspension (10^9^ CFU mL^-1^) was added to 5 mL phosphate-buffered saline and poured onto the Mueller-Hinton agar plates. After removing excess bacterial suspension by pipetting, the antibiotic discs [oxolinic acid (2 µg), florfenicol (30 µg), sulfamethoxasol/trimethoprim (25 µg) and tetracycline (30 µg)] were placed on the plates. The plates were then incubated for 3 days at 25°C. After incubation, the inhibition zone around the antibiotic discs was measured. The susceptibility patterns of the selected phage-exposed and no-phage control *F. columnare* isolates to the antibiotics were compared to that of the parent wild-type isolates.

### Motility/Colony spreading

The effect of phage-exposure on bacterial motility was tested by comparing the colony spreading ability of phage-exposed and no-phage control isolates with that of their parent wild-type isolates. After spotting of 5 μL of bacterial suspension (10^9^ CFU mL^-1^) on TYES agar (0.5% agar) plates supplemented with 0.1% baker’s yeast and incubation for 3 days at 25°C, the colony diameter of each isolate was measured. Each isolate was tested in three replicates.

### Adhesion and biofilm formation

Changes in adherence or biofilm formation capacities between wild-type, phage-exposed and no-phage control *F. columnare* isolates were studied in flat-bottomed 96-well microtiter plates (Nunclon Δ Surface, Nunc) (51). *F. columnare* cells grown on TYES agar were suspended in autoclaved fresh water (lake Littoistenjärvi) to a concentration of 10^8^ CFU mL^-1^ (OD_520nm_=0.6). For testing of bacterial adherence, a 100 μL volume of the prepared bacterial suspensions were added in triplicate into wells of replicate microtiter plates and incubated statically for 1 h at 25°C. For testing of biofilm formation, a 100 μL volume of TYES broth was added to wells containing 100 μL of the prepared bacterial suspensions and allowed to incubate for 3 days. Autoclaved fresh water was used as negative control. After incubation, the contents were discarded and the wells were washed three times with sterile 0.5% NaCl to remove non-adherent cells and air dried. The wells were then stained with 0.1% crystal violet solution for 45 min and washed three times by submersion in a container of tap water and air dried. The crystal violet was solubilized with 96% ethanol for 15 min before measuring the absorbance (1 s) at 595 nm (Victor2, Wallac).

### Protease activity

Changes in protease activity was examined by spotting 1 μL of bacterial TYES broth suspension (10^8^ CFU mL^-1^) of the wild-type isolates and each phage-exposed and no-phage control isolate on TYES agar (1.5% agar) supplemented with (w/v) elastin (0.1%), gelatin (3%) and skim milk (5%) (caseinase production). The proteolytic activity of each isolate was observed by the presence of a clear zone surrounding the colony after incubation, and assessed by measuring the clear zone ratio (diameter of clear zone/diameter of the colony) of three replicate samples. In the absence of a clearing zone outside the colony, the clear zone ratio was defined as 1. The measurements were made after 5 (caseinase and gelatinase) or 10 days (elastinase) of incubation at 25°C.

### Virulence

Virulence of phage-exposed and no-phage control *F. columnare* isolates was tested on 1.94 g (average weight) rainbow trout fry and compared to the virulence of wild-type isolates. Fifteen fish per treatment, 20 in control treatment with no bacteria, were exposed individually in 500 mL of bore hole water (25°C) to cells of single bacterial isolates by constant immersion (5.0 × 10^3^ CFU mL^-1^). Survival of the fish was monitored hourly during 24 h. Morbid fish that did not respond to stimuli were considered dead, removed from the experiment and put down by decapitation. At the end of the experiment, the fish having survived from the infection were put down using 0.008 % Benzocaine. Bacterial cultivations from gills of all the dead fish were made on Shieh agar supplemented with tobramycin (52) to confirm the presence/absence of the bacterium. Cumulative percent mortality and estimated survival time (Kaplan-Meier Survival Analysis), based on observed average survival time of fish after exposure to each isolate, were used as measures of virulence with more virulent isolates having a shorter estimated survival time.

Fish experiment was conducted according to the Finnish Act of Use of Animals for Experimental purposes, under permission ESAVI/8187/2018 granted for Lotta-Riina Sundberg by the National Animal Experiment Board at the Regional State Administrative Agency for Southern Finland.

### Whole genome sequencing

Genomes of the wild-type FCO-F2 and FCO-F9 *F. columnare* and selected (Table 1) 20 phage-exposed and four no-phage control isolates were sequenced using Illumina HiSeq platform (Institute of Molecular Medicine Finland). The Illumina data reads of FCO-F9 and its phage-exposed and no-phage control isolates were mapped to a reference genome of *F. columnare* isolate B185 (53) using Geneious software version 11.1.5 (Biomatters Ltd.). Genome of the wild-type FCO-F2 isolate was sequenced also using PacBio (BGI, China). PacBio data of FCO-F2 was assembled using > 8kbp reads with Flye (v. 2.7, four iterations) and > 6 kbp with Canu (v. 1.9). These multi-contig assemblies were then combined using Quickmerge (v. 0.3) to produce one 3 221 312 bp contig. This contig was polished with Illumina HiSeq reads using Pilon (v. 1.23), with pre-processing done using Trimmomatic (v. 0.39), bowtie2 (2.3.5.1) and Samtools (v. 1.9). The quality of the polished contig was quantified using Busco (v. 4.0.2), which reported 100% completeness of genome against the bacteria_odb10 reference set. The genome was annotated using the NCBI Prokaryotic Genome Annotation Pipeline (PGAP) (54, 55), and used as reference genome for mapping of F2 phage-exposed and no-phage control isolates.

### Statistical analyses

IBM SPSS Statistics version 24 was used for statistical analysis of the data. A one-way analysis of variance (ANOVA) was used to compare means from phenotypic analyses between experimental groups (phage-exposed isolates and no-phage control isolates) and parent wild-type isolates. If needed, lg10, exponential or square root transformations were made for the data to fulfil the homogeneity of variances assumption. If the homogeneity of variances could not be met by transformations, the data were analysed using non-parametric Kruskal-Wallis and Mann-Whitney tests. In case of elastinase and casienase activity, and biofilm formation, the isolates with no activity/biofilm forming capacity were excluded from the ANOVA LSD multiple comparison analyses. Kaplan-Meier Survival Analysis was used for analysis of virulence data.

### Data availability

The whole genome sequences of all the isolates were submitted to GenBank under accession numbers presented in Table 6.

**Table 6.** Accession numbers of whole genome sequences of wild-type *Flavobacterium columnare* isolates FCO-F2 and FCO-F9 and their phage-exposed (F2R- and F9R) and no-phage control (F2S- and F9S-) isolates submitted to GenBank.

| Isolate | Accession number |
| --- | --- |
| FCO-F2 | CPO51861 |
| F2R58 | CP054506 |
| F2R60 | CP054505 |
| F2R62 | CP054504 |
| F2R64 | CP054503 |
| F2R65 | CP054502 |
| F2R66 | CP054501 |
| F2R67 | CP054500 |
| F2R68 | CP054499 |
| F2R70 | CP054498 |
| F2R72 | CP054497 |
| F2R74 | CP054496 |
| F2S4 | CP054495 |
| F2S17 | CP054494 |
| FCO-F9 | CP054518 |
| F9R56 | CP054517 |
| F9R58 | CP054516 |
| F9R61 | CP054515 |
| F9R64 | CP054514 |
| F9R66 | CP054513 |
| F9R69 | CP054512 |
| F9R72 | CP054511 |
| F9R75 | CP054510 |
| F9R78 | CP054509 |
| F9S15 | CP054508 |
| F9S17 | CP054507 |

## Supporting information

Supplemental material: Figure S1, Table S1 and Data set S1

Supplemental Data set 1

## Acknowledgements

We acknowledge funding from Academy of Finland (grant #314939) and Jane and Aatos Erkko Foundation. This work resulted from the BONUS FLAVOPHAGE project supported by BONUS (Art 185), funded jointly by the EU and Academy of Finland.

