## Supplemental material: Figure S1, Table S1 and Data set S1 for "Bacteriophage Resistance Affects *Flavobacterium columnare* Virulence Partly via Mutations in Genes Related to Gliding Motility and Type IX Secretion System"

[1]Department of Biological and Environmental Science and Nanoscience Center, University of Jyväskylä, Jyväskylä, Finland; [2]Laboratory of Aquatic Pathobiology, Åbo Akademi University, Turku, Finland; [3]Department of Biology, Marine Biological Section, University of Copenhagen, Helsingør, Denmark; *Present address: Department of Biological Sciences, University of Bergen, Bergen, Norway

### Supplemental material

The phage-exposed and no-phage control isolates showed antibiotic susceptibility patterns similar to the parent wild-type isolates (Figure S1). Most showed decreased inhibition zone diameter in trimethoprim/sulfamethoxazole test, but including wild-type isolates, this inhibition zone was weak and not totally clear from bacterial growth. In case of other antibiotics, decrease in inhibition zone diameter was within a range of measurement error (1-2 mm) (Table S1). However, these results are based on only one repeat so statistical analyses could not be conducted and no estimates on significance of the results can be made.

In addition to statistical differences between the virulence of phage-sensitive rhizoid and phage-resistant rough and soft isolates, there were also differences between the virulence of individual isolates (Data set S1).

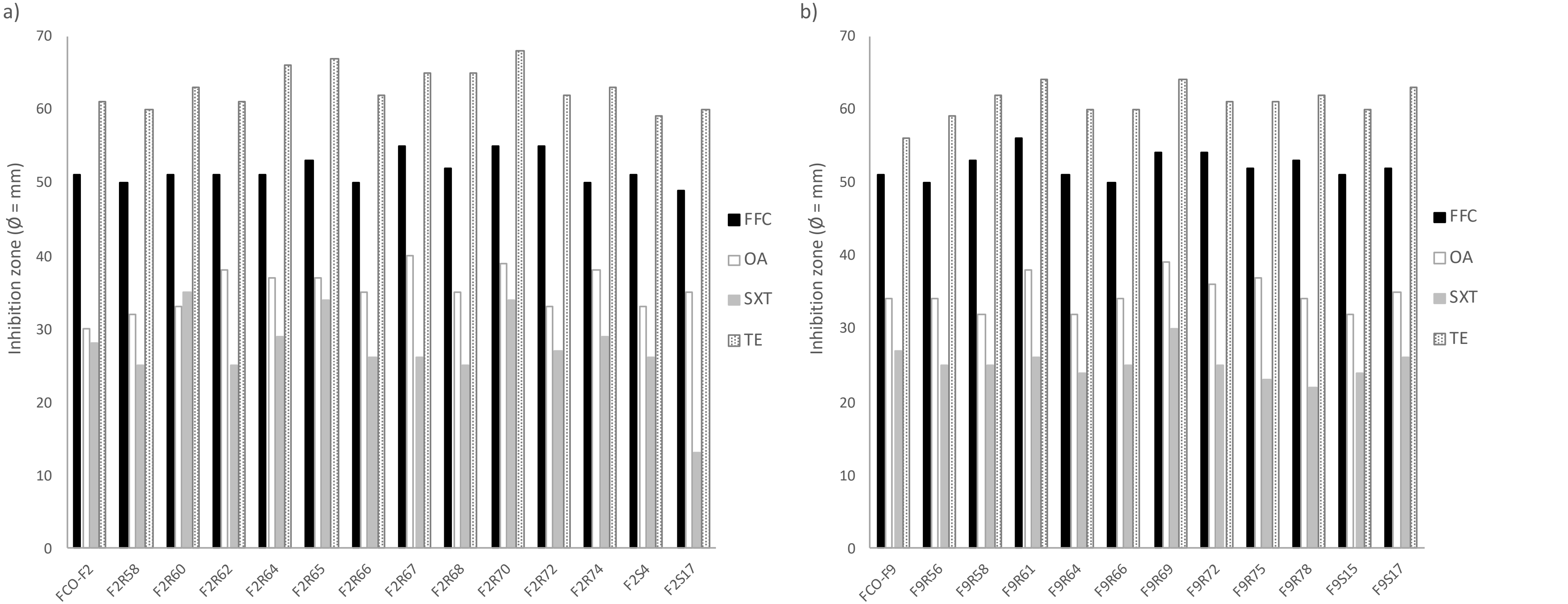


**Figure S1.** Antibiotic susceptibility of the wild-type *Flavobacterium columnare* a) FCO-F2 and b) FCO-F9 isolates, and their phage-exposed (F2R- and F9R-) and no-phage control (F2S- and F9S-) isolates against florfenicol (FFC), oxolinic acid (OA), sulfamethoxasol/trimethoprim (SXT) and tetracycline (TE) measured as the inhibition zone diameter (mm) with the Kirby-Bauer disc diffusion method.

**Table S1.** Antibiotic susceptibility of the wild-type *Flavobacterium columnare* FCO-F2 and FCO-F9 isolates, and their phage-exposed (F2R- and F9R-) and no-phage control (F2S- and F9S-) isolates against florfenicol (FFC), oxolinic acid (OXA), sulfamethoxasol/trimethoprim (SXT) and tetracycline (TE) measured as the inhibition zone diameter (mm) with the Kirby-Bauer disc diffusion method. Values of the wild-type isolates are underlined, and values with bold indicate decreased inhibition zone diameter compared to the parent wild-type isolate.

| Isolate | Inhibition zone diameter (mm) | | | |
|---|---|---|---|---|
| | FFC | OA | SXT | TE |
| FCO-F2 | 51 | 30 | 28 | 61 |
| F2R58 | **50** | 32 | **25** | **60** |
| F2R60 | 51 | 33 | 35 | 63 |
| F2R62 | 51 | 38 | **25** | 61 |
| F2R64 | 51 | 37 | 29 | 66 |
| F2R65 | 53 | 37 | 34 | 67 |
| F2R66 | **50** | 35 | **26** | 62 |
| F2R67 | 55 | 40 | **26** | 65 |
| F2R68 | 52 | 35 | **25** | 65 |
| F2R70 | 55 | 39 | 34 | 68 |
| F2R72 | 55 | 33 | **27** | 62 |
| F2R74 | **50** | 38 | 29 | 63 |
| F2S4 | 51 | 33 | **26** | **59** |
| F2S17 | **49** | 35 | **13** | **60** |
| FCO-F9 | 51 | 34 | 27 | 56 |
| F9R56 | **50** | 34 | **25** | 59 |
| F9R58 | 53 | **32** | **25** | 62 |
| F9R61 | 56 | 38 | **26** | 64 |
| F9R64 | 51 | **32** | **24** | 60 |
| F9R66 | **50** | 34 | **25** | 60 |
| F9R69 | 54 | 39 | 30 | 64 |
| F9R72 | 54 | 36 | **25** | 61 |
| F9R75 | 52 | 37 | **23** | 61 |
| F9R78 | 53 | 34 | **22** | 62 |
| F9S15 | 51 | **32** | **24** | 60 |
| F9S17 | 52 | 35 | **26** | 63 |

**Data set S1.** Statistical differences (Kaplan-Meier Survival Analysis, pairwise comparisons; Log-Rank, Mantel Cox) between cumulative mortalities of rainbow trout caused by wild-type *Flavobacterium columnare* FCO-F2 and FCO-F9 isolates, and their phage-exposed (F2R- and F9R-) and no-phage control (F2S- and F9S-) isolates.

(SEPARATE EXCEL FILE)
